# Taurine Transporter SLC6A6 Expression Promotes Mesenchymal Stromal Cell Function

**DOI:** 10.1101/2025.06.24.661367

**Authors:** Christina M. Kaszuba, Sonali Sharma, Benjamin J. Rodems, Cameron D. Baker, Palomi Schacht, Takashi Ito, Kyle P. Jerreld, Chen Yu, Edgardo I. Franco, Emily R. Quarato, Francisco A. Chaves, Jane L. Liesveld, Laura M. Calvi, Hani A. Awad, Roman A. Eliseev, Jeevisha Bajaj

## Abstract

Mesenchymal stromal cell (MSC) differentiation is critical for the development, maintenance, and repair of bone tissue. This occurs within the bone marrow microenvironment, which consists of various stromal populations and structural components that support skeletal growth, repair and homeostasis. MSCs also promote self-renewal of hematopoietic stem cells and regulate their differentiation. Analysis of our own and publicly available single-cell RNA sequencing datasets of the murine non-immune bone and bone marrow indicates that Slc6a6 expression is enriched in MSCs. Slc6a6 encodes for an ion-dependent transporter of taurine (TauT). Although taurine supplements have been shown to mitigate the onset of bone defects in aged populations, the absence of TauT expression in osteolineage cells suggests that taurine’s effect on bone may be secondary to its function in other populations, such as MSCs. Using young TauT genetic loss-of-function murine models we find that TauT loss impacts MSC populations *in vivo* and impairs MSC osteogenic differentiation *in vitro*. This correlates with decreased bone mineral density and bone strength in young TauT knockout mice. Importantly, shRNA-based knockdown of SLC6A6 expression in primary human MSCs reduces osteogenic differentiation, indicating a key role of taurine uptake in human MSC function. Consistent with a decline in MSC function with TauT loss, we find that TauT null MSCs are unable to support self-renewal and expansion of co-cultured hematopoietic stem/progenitor populations. Mechanistically, our RNA-sequencing analysis identifies downregulation of Wnt/beta-catenin signaling in MSCs in the absence of TauT. These cells also show reduced oxidative phosphorylation, and increased ROS levels, indicating that impaired Wnt signaling and elevated oxidative stress may contribute to the observed defects in osteogenic differentiation capacity. Collectively, our data identify taurine uptake as a key regulator of mesenchymal stromal cell maintenance and osteogenic fate determination.

## INTRODUCTION

The bone marrow microenvironment (BMME) is a complex system consisting of hematopoietic cells, non-hematopoietic stromal populations, and extracellular matrix. The BMME plays a critical role in providing mechanical and structural support to bone, promoting self-renewal of stromal populations, and regulating lineage differentiation (1, 2). Mesenchymal stromal cells (MSCs), are one of the key cell types in the BMME and play a pivotal role in bone function due to their ability to differentiate into cells critical for bone development including osteoblasts, adipocytes, fibroblasts, and chondrocytes (3). MSCs are also essential for the development and support of hematopoietic stem/progenitor cells, which are responsible for the production and maintenance of all blood and immune cell types (4). Despite their importance for multiple physiological functions, molecular mechanisms that sustain MSC function and maintenance are not fully understood. Elucidating mechanisms that promote MSC fitness may help inform therapies promoting bone health and hematological support in the long term.

MSC differentiation into osteoblasts promotes bone formation through matrix mineralization (5). Osteoblasts can terminally differentiate into osteocytes, the most abundant cell skeletal cell, which maintain the bone matrix (6). Bone undergoes constant remodeling, where osteoclasts resorb old bone and osteoblasts form new bone (7). Defective MSC differentiation and mineralization have been linked to bone disorders like osteoporosis/osteopenia, osteomalacia, and osteogenesis imperfecta, leading to compromised bone integrity, fragility, and reduced bone mass (8–10). There is thus a critical need to better identify mechanisms that can promote MSC function to develop therapeutic interventions for millions of people that suffer from osteoporosis.

MSC differentiation along the osteolineage pathway is stimulated by growth factors such as transforming growth factor-beta (TGF-β), bone morphogenic proteins (BMPs) (11), fibroblast growth factor (FGF) (12), transcription factors including Osterix, Runx2, and β-catenin (13, 14), and signaling pathways including hedgehog (15), Notch (16), and wingless/integrated (Wnt) (17, 18). In addition to signaling molecules, metabolic changes within the BMME are known to play a key role in bone homeostasis, and can influence bone formation and remodeling, in part by regulating MSCs. *In vivo* and *in vitro* studies identified that glutamine consumption is critical for the function and differentiation MSCs along both the osteolineage and adipolineage pathway, and necessary to maintain bone mineral density (19, 20). Arginine and lysine are key components of collagen, which are critical for bone formation and structure (19, 21). Osteoblasts cultured with exogenous arginine and lysine show significant increases in type 1 collagen synthesis, improved cell viability, and prevention of senescence (21). Recent work indicates that the non-essential amino acid taurine can inhibit age related bone loss in 24-month-old mice, indicating an important role of taurine in promoting bone remodeling and bone health in the context of aging (22–24). However, it is not known if taurine directly impacts mature bone cells in young and old populations.

Taurine is present in high amounts in mammalian tissues like the brain, heart, retina, and skeletal system (23, 25). While diet is the primary source of taurine, with major amounts found in meat and fish, it can also be synthesized *in vivo* from cysteine in cells expressing the enzymes cysteine dioxygenase (CDO1) and cysteine sulfinic acid decarboxylase (CSAD) (11). Taurine exerts multiple physiological functions including serving as an organic osmolyte, cytoprotection, and antioxidant defense (26–29). In addition, taurine uptake can promote glycolysis and thus fuel the growth of aggressive myeloid leukemia cells (30). These functions rely on intracellular taurine levels, which are primarily controlled by the expression of the taurine transporter, TauT. TauT, encoded by the SLC6A6 gene, is the primary, high affinity sodium chloride-dependent taurine transporter (Km < 10 µM), with a much lower affinity for β-alanine (Km = 55 µM) (30).

In human osteoblast and osteocyte cultures *in vitro*, taurine supplementation has been shown to increase cell proliferation and protect against oxidative stress-induced cell death (29, 31, 32). However, the role of taurine in regulating the differentiation fate of MSCs, especially in the context of young populations, remains unclear. Here, we use *Slc6a6* knockout mice to determine the impact of loss of taurine uptake on MSC function in young, 16-week-old animals. Our work identifies the importance of taurine uptake on MSC function and bone health, providing further insight into its role in MSC maintenance, regenerative therapies, and treatment for bone diseases.

## RESULTS

### Mesenchymal stromal cells have the highest expression of Slc6a6

While taurine supplements have been shown to promote bone health, it is not known if taurine is directly taken up by the osteoblasts or if the impact on bone health is an indirect effect of taurine uptake by other bone marrow populations. To test this, we analyzed taurine transporter (Slc6a6) expression in single-cell RNA-sequencing (scRNA-seq) datasets of non-immune murine bone and bone marrow populations in 8-week old mice in our own dataset (30), as well as other publicly available scRNA-seq datasets of 6-22 week old mice (33, 34, 35). Our analysis indicated that while all MSCs and a small subset of endothelial cells had high Slc6a6 expression (Fig. 1a, b, Fig. S1a), it was not detected in cells differentiating along the osteolineage pathway. Further, MSCs did not express other transporters that have been shown to non-specifically transport taurine at much lower efficiencies, including Slc6a13, Slc16a6, Slc36a1 (36–38) (Fig. S1b). Consistent with the scRNA-seq analysis, our experiments show that MSCs undergoing osteogenic differentiation rapidly downregulate Slc6a6 expression and have a 30-fold increase in Bglap expression, a common marker of osteogenic differentiation (Fig. 1c-f). Since MSCs can also differentiate along the adipogenic lineage, we tested the impact of adipogenic differentiation on Slc6a6 expression. Our experiments showed that similar to osteolineage cells, adipocytes have 15-fold lower Slc6a6 expression and 5-fold higher Pparg expression, a common marker of adipogenic differentiation, when compared to undifferentiated MSCs (Fig. 1g-j). These data indicate that Slc6a6 is primarily expressed in undifferentiated MSCs. We thus focused our studies on MSCs as they are known to be critical for bone development and maintenance.

**Figure 1:**
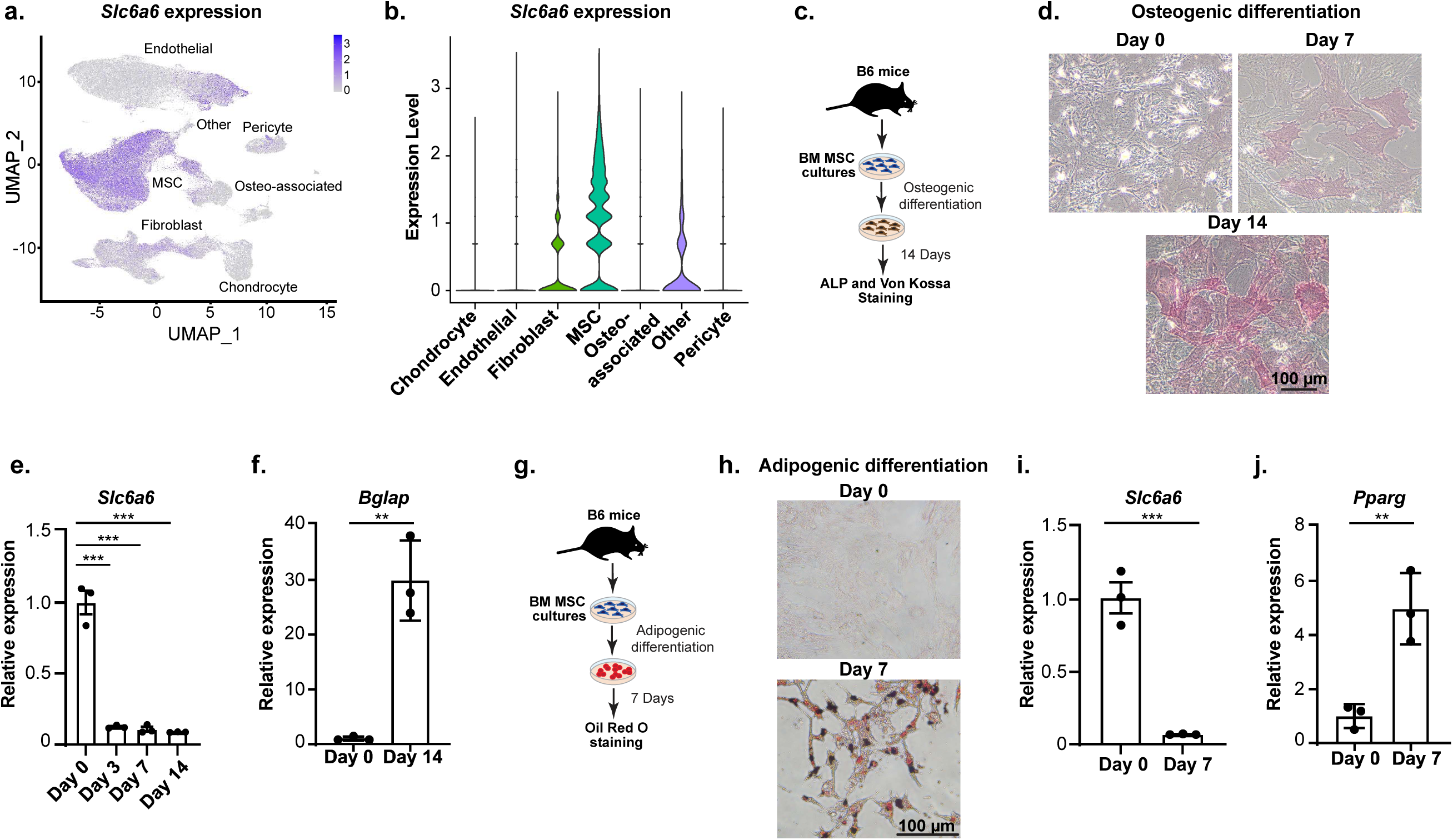
Mesenchymal stromal cells have high Slc6a6 expression. **a,** Slc6a6 expression in 7 distinct non-hematopoietic bone and bone marrow stromal cell clusters (scale bar represents expression level). **b,** Violin plot of Slc6a6 expression in non-hematopoietic cells (Data from GSE226644, GSE108892, GSE122467, GSE128423). **c**, Experimental strategy for MSC osteogenic differentiation. **d-f,** Microscopy images of alkaline phosphatase and Von Kossa staining (**d**), Slc6a6 expression (**e**), and Bglap expression (**f**) in MSCs undergoing osteogenic differentiation (**e**, **f**, mean ± s.d.; n=3 technical replicates per cohort; **e**, one-way ANOVA). **g,** Experimental strategy for MSC adipogenic differentiation. **h-j** Oil Red O staining (**h**), Slc6a6 expression (**i**), and Pparg expression (**j**) in MSCs undergoing adipogenic differentiation (**i**, **j**, mean ± s.d.; n=3 technical replicates per cohort) (*p<0.05, **p<0.01. ***p<0.001). All analyses are from unpaired two-tailed Student’s t-test or as indicated.

### Impact of TauT loss on MSC and endothelial populations *in vivo*

The high expression of Slc6a6 (TauT) in MSCs and some endothelial populations indicates that taurine uptake may be essential for sustaining these populations. To test this, we used a global TauT knockout murine model (39). While TauT knockout mice are born in Mendelian ratios, they can develop functional impairments in the skeletal system and eye with age (23, 39, 40). Our LC/MS based assays identified a profound 231-fold reduction in taurine levels in MSCs from TauT^-/-^ mice as compared to wild type controls (Fig. 2a, b), confirming that TauT mediated taurine uptake is the primary source of taurine in these cells.

**Figure 2:**
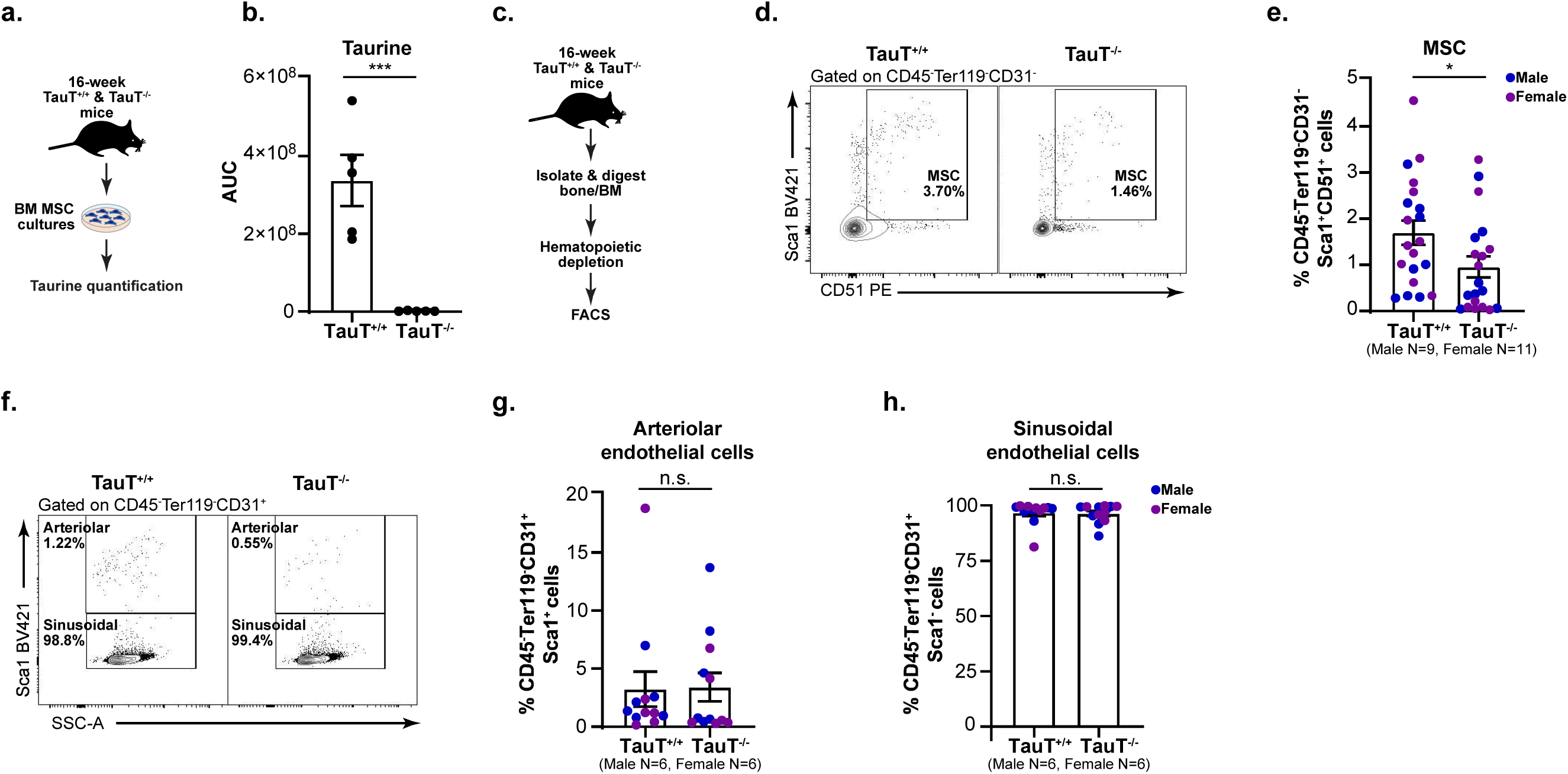
Impact of TauT loss on bone marrow microenvironmental populations *in vivo*. **a**, **b**, Experimental strategy (**a**) and taurine quantification (**b**) in MSCs (mean ± s.e.m.; n=5 independent cultures per cohort). **c,** Experimental strategy used to analyze microenvironmental cells in TauT^+/+^ and TauT^-/-^ mice. **d**, **e**, Representative FACS plots (**d**) and quantification of MSCs (**e**) in bone marrow stroma (mean ± s.e.m.; data are combined from seven independent experiments). **f-h**, Representative FACS plots (**f**) and quantification of arteriolar endothelial cells (**g**) and sinusoidal endothelial cells (**h**) in bone marrow stroma (mean ± s.e.m.; data are combined from five independent experiments) (*p<0.05, **p<0.01. ***p<0.001). All analyses are from unpaired two-tailed Student’s t-test or as indicated.

To determine if TauT loss impacts bone marrow microenvironmental populations *in vivo*, we used flow cytometry to analyze the non-hematopoietic stromal populations in the femur of TauT^-/-^ and TauT^+/+^ mice (Fig. 2c). Our experiments showed a 44% decrease in MSC frequency with TauT loss. However, we could not detect any difference in either arteriolar or sinusoidal endothelial cells (Fig. 2d-h). Our flow cytometry experiments, in combination with our analysis of scRNA-seq data, indicate that while TauT expression is essential for sustaining MSC populations in the bone marrow at a steady state, it may be dispensable for maintaining endothelial subsets.

### Functional effect of TauT loss on MSC function *in vitro*

Our experiments indicate that taurine may be important for sustaining MSC populations within the bone marrow. We thus determined the functional impact of TauT loss on MSCs. MSCs can differentiate into multiple cell types including osteoblast, adipocytes, fibroblasts, and chondrocytes (3). To test if TauT loss impairs proliferation and differentiation of MSCs, we tested the fibroblast (CFU-F), and osteogenic differentiation (CFU-OB) capacity of bone marrow cells derived from TauT^+/+^ and TauT^-/-^ mice (Fig. 3a). Our experiments identified a 41-52% decrease in the CFU-F forming ability (Fig. 3b, c) of cells isolated from TauT^-/-^ mice as compared to wild-type controls, indicating reduced proliferation capacity. In addition, TauT^-/-^ cells formed 62% fewer CFU-OBs (Fig. 3d, e) when compared to wild-type controls, indicating impaired osteogenic differentiation potential.

**Figure 3:**
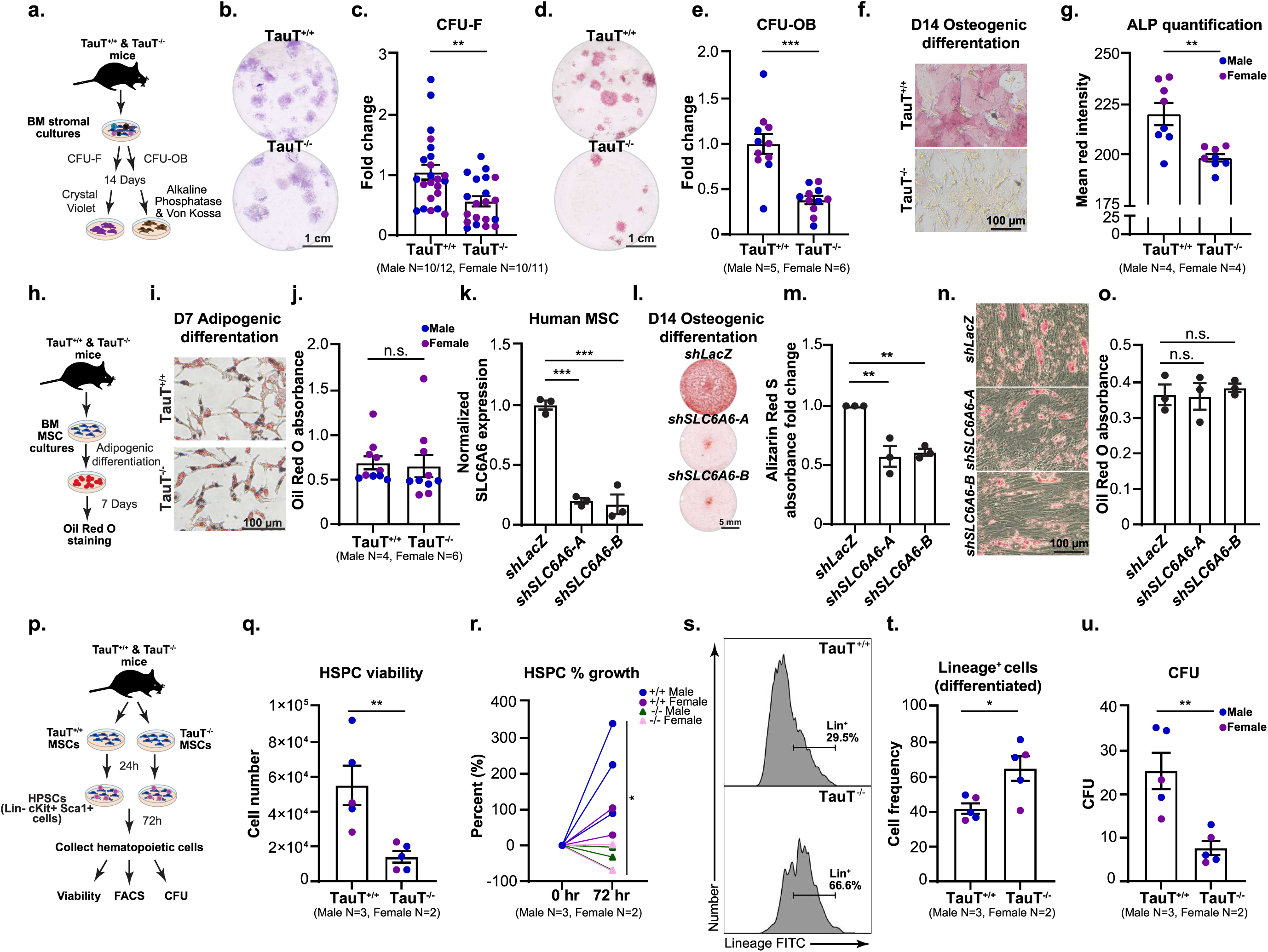
Functional effect of TauT loss on MSC function *in vitro*. **a,** Experimental strategy used for colony forming unit fibroblast (CFU-F) and osteoblast (CFU-OB) assays from bone marrow cells from TauT^+/+^ and TauT^-/-^ mice. **b**, **c**, Crystal Violet staining (**b**) and relative CFU-F quantification (**c**) (mean ± s.e.m.; data are combined from four independent experiments). **d**, **e**, Alkaline phosphatase (ALP) and Von Kossa staining (**d**) and relative CFU-OB quantification (**e**) (mean ± s.e.m.; data are combined from four independent experiments). **f**, **g,** Microscopy images of ALP and Von Kossa staining (**f**) and quantification (**g**) in MSCs undergoing osteogenic differentiation for 14 days (mean ± s.e.m.; data are combined from three independent experiments). **h,** Experimental strategy for MSC adipogenic differentiation. **i**, **j**, Oil Red O staining (**i**) and absorbance quantification (**j**) in MSCs undergoing adipogenic differentiation (mean ± s.e.m.; data are combined from three independent experiments). **k,** Normalized SLC6A6 expression in primary human donor bone marrow derived MSCs transduced with shRNAs targeting *LacZ* (control) or *SLC6A6* (mean ±s.d.; n=3 technical replicates per cohort; one-way ANOVA). **l**, **m,** Alizarin Red S staining (**l**) and quantification (**m**) in primary human donor bone marrow derived MSCs undergoing osteogenic differentiation for 14 days (mean ±s.e.m.; data are combined from three independent experiments; one-way ANOVA). **n**, **o**, Day 7 Oil Red O staining (**n**) and absorbance quantification (**o**) in primary human bone marrow derived MSCs transduced with shRNAs targeting LacZ (control) or SLC6A6 undergoing adipogenic differentiation (mean ± s.d.; n=3 independent culture wells per cohort; one way ANOVA). **p**, Experimental strategy used for MSC and hematopoietic stem/progenitor cell (HSPC) co-culture. **q**, Trypan blue based cell viability of HSPCs 72 hours post coculture with TauT^+/+^ and TauT^-/-^ MSCs (mean ±s.e.m.; data are combined from four independent experiments). **r**, Percent growth of HSPCs 72 hours post coculture with TauT^+/+^ and TauT^-/-^ MSCs (mean ±s.e.m.; data are combined from four independent experiments; shape represents genotype). **s**, **t**, Representative FACS histogram (**s**) and quantification (**t**) of Lineage^+^ frequency of HSPCs 72 hours post coculture with TauT^+/+^ and TauT^-/-^ MSCs (mean ±s.e.m.; data are combined from four independent experiments). **u**, CFU of HSPCs 72 hours post coculture with TauT^+/+^ and TauT^-/-^ MSCs (mean ±s.e.m.; data are combined from four independent experiments) (*p<0.05, **p<0.01. ***p<0.001). All analyses are from unpaired two-tailed Student’s t-test or as indicated.

To confirm that MSCs, and no other cell types in the bone marrow, contribute to the observed CFU defects in TauT^-/-^ mice, we established purified bone marrow MSC cultures from TauT^-/-^ and TauT^+/+^ mice using established protocols (41). We then tested their ability to undergo osteogenic and adipogenic differentiation over 7-14 days. Our quantification of alkaline phosphatase (ALP) and Von Kossa-stained cells 14 days post induction of differentiation identified a significant defect in the ability of TauT^-/-^ MSCs to undergo osteogenic differentiation (Fig. 3f, g). However, we did not identify any defects in the ability of TauT^-/-^ MSCs to differentiate along the adipogenic lineage (Fig. 3h-j). These data suggest that while taurine uptake by MSCs is essential for determination of osteogenic fate, it may not impair the adipogenic potential of murine MSCs.

Next, we confirmed if the observed defects with TauT loss in murine MSCs are also relevant to human cells. To this end, we used two independent shRNAs to knockdown SLC6A6 expression in primary human donor-derived bone marrow MSCs by 5-fold (Fig. 3k). We then induced osteogenic differentiation of these human MSCs transduced with shRNAs targeting *SLC6A6* or *LacZ* for 14 days. Consistent with our murine data, human MSCs lacking *SLC6A6* expression showed 39-42% lower ability to undergo osteogenic differentiation as compared to controls (Fig. 3l, m). However, loss of SLC6A6 expression in human MSCs did not impact adipogenic differentiation (Fig. 3n,o).

Thus, our experiments show that TauT loss leads to functional defects in MSC proliferation and osteogenic fate, which may lead to weaker bones *in vivo*.

### TauT loss impairs MSCs ability to support hematopoietic stem and progenitor cells *in vitro*

In addition to MSC function in bone formation, they are also known to support hematopoietic stem/progenitor cells (HSPCs) *in vivo* (42, 43). MSCs can secrete growth factors (SCF, VEGF, FGF2, HGF), chemokines (CCL2, CXCL12), and cytokines (IL-10) that are known to support HSPC proliferation and survival (44, 45). Our experiments thus far identify that TauT loss impairs MSC osteogenic fate. We thus tested if loss of TauT also impairs the ability of MSCs to support HSPCs *in vitro*. To this end, we carried out co-culture assays of TauT^+/+^ and TauT^-/-^ MSCs with wild-type HSPCs (CD3^-^CD4^-^CD8^-^CD19^-^B220^-^CD11^-^Ly6G^-^/Ly6C^-^Ter119^-^cKit^+^Sca1^+^; Fig. 3p). Our experiments show that HSPCs co-cultured with TauT^-/-^ MSCs for 72 hours had 4-fold reduced viability (Fig 3q) and a 4.5-fold lower percent growth (Fig. 3r). This loss in growth and viability was accompanied by a 40% increase in frequency of Lin^+^ differentiated hematopoietic cells (Fig. 3s, t). In addition, the functional ability of co-cultured HSPCs to form colonies in methylcellulose was reduced by 3-fold (Fig. 3u). These data indicate that MSCs from TauT^-/-^ mice have functional defects that impair their ability to support the expansion of HSPCs *in vitro*.

### Physical and mechanical properties of TauT^-/-^ murine model *in vivo*

MSCs are known to be critical for bone formation and maintenance. Our experiments thus far have established that TauT loss leads to a reduction in MSCs at steady state, functional defects in MSC proliferation and osteogenic fate determination, and impaired ability of MSCs to support HSPCs. We thus investigated the effect of TauT loss on physical *in vivo* bone properties in both male and female young adult mice at 16-weeks of age. We first used dual-energy X-ray absorptiometry (DEXA) scanning to evaluate femurs and whole-body composition in our TauT^-/-^ murine model for 38 weeks. The weight of each mouse was recorded prior to each DEXA scan and showed that TauT^-/-^ mice had 12-19% lower weight than wild-type controls at 16-weeks (Fig. 4a). The DEXA scans also revealed a 14% decrease in femur bone mineral density in female TauT^-/-^ mice beginning at 16-weeks, a time point consistent with adult characteristics (46) (Fig. 4b). However, there was no change in femur fat percentage (Fig. S2a), consistent with our *in vitro* data showing that TauT loss does not impact MSC adipogenic differentiation (Fig. 3i, j).

**Figure 4:**
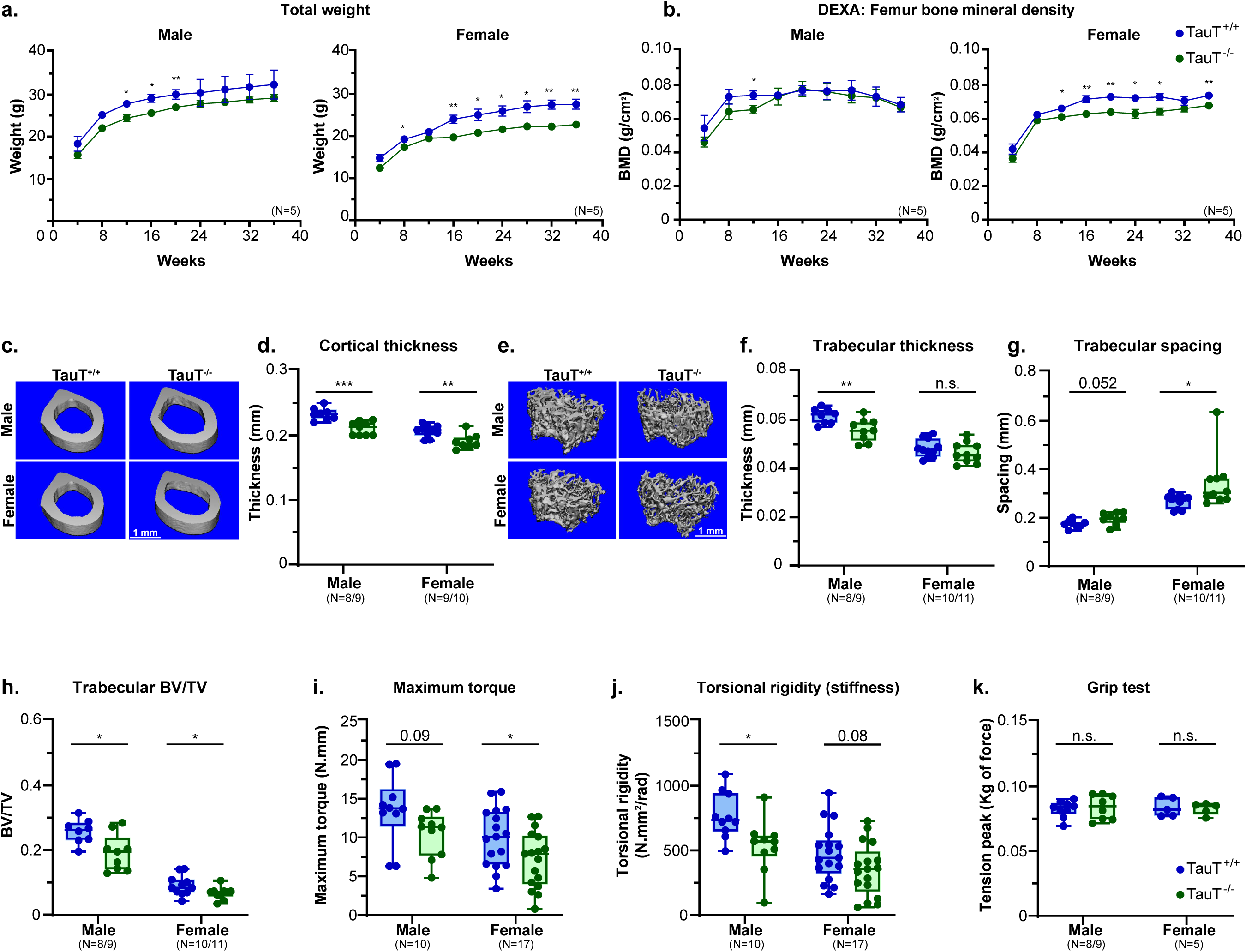
Physical and mechanical properties of TauT^-/-^ murine model *in vivo*. **a**, Total weight of TauT^+/+^ and TauT^-/-^ mice over a 40-week period (mean ±s.e.m.; data are combined from four independent experiments). **b**, Dual x-ray absorptiometry (DEXA) scans of femur bone mineral density (BMD) of TauT^+/+^ and TauT^-/-^ mice over a 40-week period (mean ±s.e.m.; data are combined from four independent experiments). **c**, **d**, Micro-CT pictographs of femur cortical region (**c**) and quantification of cortical thickness (**d**) (mean ±s.e.m.; data are combined from three independent experiments). **e-h**, Micro-CT pictographs of femur trabecular region (**e**), quantification of trabecular thickness (**f**), trabecular spacing (**g**), and trabecular bone volume/total volume (**h**) (mean ±s.e.m.; data are combined from three independent experiments). **i**, **j**, Biomechanical torsion testing of tibia from TauT^+/+^ and TauT^-/-^ mice, quantification of maximum torque (**i**) and torsional rigidity (**j**) (mean ±s.e.m.; data are combined from three independent experiments). **k**, Grip strength testing in TauT^+/+^ and TauT^-/-^ mice (mean ±s.e.m.; data are combined from five independent experiments) (*p<0.05, **p<0.01. ***p<0.001). All analyses are from unpaired two-tailed Student’s t-test.

Our analysis of female and male TauT^-/-^ femur bone morphological structure using micro-computed tomography (micro-CT) revealed a 5-10% decrease in both cortical and trabecular thickness, a 11-21% increase in trabecular spacing, and a ∼30% decrease in trabecular BV/TV when compared to TauT^+/+^ controls, indicating an osteopenia-like (47) phenotype (Fig. 4c-h). However, we noted no difference in tartrate-resistant acid phosphatase (TRAP) staining of osteoclasts in TauT^-/-^ murine femurs when compared to TauT^+/+^ controls (Fig. S2b), indicating that decreased physical bone properties likely reflect defects in MSC function, and not in osteoclasts.

To further determine how bone structural changes affect the biomechanical properties in TauT^-/-^ mice, we carried out torsion testing in tibias. Our torsion analysis showed that both male and female TauT^-/-^ tibias had a 22-28% decrease in maximum torque and a 28-31% reduction in torsional rigidity (Fig. 4i, j). This biomechanical testing suggests that TauT^-/-^ mice have weaker, more brittle bones when compared to wildtype controls. Finally, since taurine can be found in the muscle (48), we also evaluated grip strength in the TauT^-/-^ murine model. We noted no differences in grip strength with TauT loss in 16-week-old mice (Fig. 4k). Moreover, our histological analysis on murine gastrocnemius muscles showed no changes in muscle fiber/nuclei structure or cross-sectional area (Fig. S2c), indicating that the observed defects in bone quality are not an indirect effect of altered muscle function in young adult mice.

Collectively, our data identify a decline in the *in vivo* bone physical and mechanical properties with TauT loss in young-adult mice.

### Decreased Wnt/beta-catenin signaling and oxidative phosphorylation contribute to MSC osteogenic defects in the absence of TauT

To determine the underlying mechanisms by which taurine loss alters MSC functional properties *in vitro* and *in vivo*, we carried out RNA-sequencing based gene-expression analysis in male and female TauT^+/+^ and TauT^-/-^ MSCs (Fig. S3a). Our RNA-sequencing identified 1,139 downregulated and 951 upregulated genes in the absence of TauT (Padj<0.05) (Fig. 5a).

**Figure 5:**
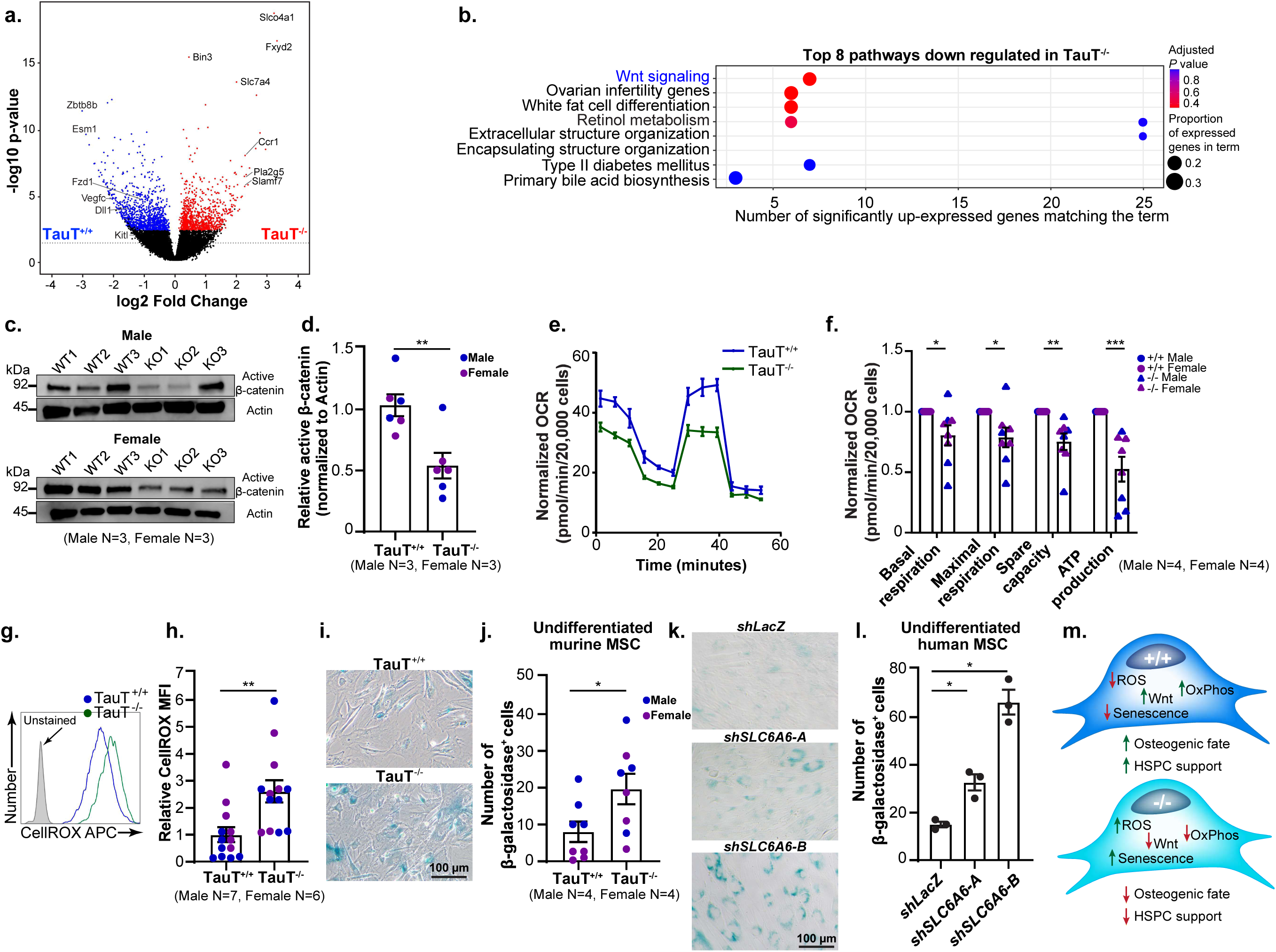
Loss of TauT impairs Wnt/beta-catenin signaling and oxidative phosphorylation in MSCs **a**, Volcano plot of genes upregulated in TauT^+/+^ (blue) and TauT^-/-^ (red) MSCs (n=8 male and n=7 female MSCs). **b**, Gene sets significantly downregulated in TauT^-/-^ MSCs (n=8 male and n=7 female MSCs; blue text indicates pathways of interest). **c**, **d,** immunoblot (**c**), and densitometric quantification (**d**) of indicated proteins (mean ±s.e.m.). Original western blots are in Supplemental Material. **e**, **f**, Curve of oxygen consumption rate (OCR) (**e**) and normalized quantification (**f**) (mean[±[s.e.m.; n=4-5 independent culture wells per cohort; data combined from four independent experiments). **g**, **h**, Representative CellROX histogram (**g**) and quantification of mean fluorescent intensity in TauT^+/+^ and TauT^-/-^ MSCs (**h**) (mean ±s.e.m.; data are combined from four independent experiments). **i**, **j**, Microscopy images of senescence-associated β-galactosidase staining (SA-β-gal) (blue) (**i**) and quantification (**j**) of SA-β-gal in undifferentiated murine MSCs (mean[±[s.e.m.; data combined from three independent experiments). **k**, **l**, Microscopy image of SA-β-gal (blue) (**k**) and quantification (**l**) of SA-β-gal in undifferentiated primary human donor bone marrow derived MSCs transduced with shRNAs targeting *SLC6A6* or *LacZ* (mean[±[s.e.m.; n=3 independent culture wells from one primary human donor bone marrow derived MSC sample; one-way ANOVA). **m**, Schematic depicts how taurine regulates MSC function (*p<0.05, **p<0.01. ***p<0.001). All analyses are from unpaired two-tailed Student’s t-test or as indicated.

Enrichr analysis revealed that the downregulated genes in TauT^-/-^ MSCs primarily constituted pathways associated with the canonical Wnt/β-catenin signaling, and included Wnt-pathway related genes such as Esm1, Fzd1, Zbtb8b, and Dll1 (Fig. 5a, b). The upregulated genes in TauT^-/-^ MSCs primarily constituted pathways associated with inflammation and immune response, and included genes such as Bin3, Ccr1, and Slamf7 which are known to regulate pro-inflammatory responses (Fig. S3b and Fig. 5b).

In murine models, Wnt/beta-catenin signaling has been associated with low bone mass (49, 50). To confirm our RNA-Seq based analysis, we determined levels of active β-catenin in TauT^+/+^ and TauT^-/-^ MSCs. Our Western-blot assays showed that TauT loss results in nearly a 48% reduction in non-phosphorylated active beta-catenin (β-catenin) as compared to controls (Fig. 5c, d), confirming that Wnt signaling is downregulated with TauT loss in MSCs. To identify mechanisms that may lead to Wnt pathway downregulation, we tested if oxidative phosphorylation is affected since it is known to regulate canonical Wnt/beta-catenin signaling (17). We thus evaluated extracellular metabolism of TauT^+/+^ and TauT^-/-^ MSCs in vitro by measuring oxygen consumption rate (OCR) and extracellular acidification rate (ECAR). Our experiments showed that TauT^-/-^ MSCs had a ∼20% decrease in basal respiration, maximal respiration, and in spare capacity, and a 47% decrease in ATP production as compared to wild-type controls (Fig. 5e, f). However, we did not identify major differences in glycolysis in the absence of TauT (Fig. S3c, d).

Oxidative stress is a result of the imbalance between production of reactive oxygen species (ROS) and the cell’s ability to remove ROS. Further, it is known that ROS play an important role in immune inflammatory response (51, 52). Elevated levels of reactive oxygen species (ROS) are known to impair Wnt/Beta-catenin signaling (53). Taurine functions as an antioxidant and protects cells against oxidative stress induced cell damage (54). Consistent with the downregulation of Wnt/beta-catenin signaling and the upregulation of pathways associated with immune response in TauT^-/-^ MSCs (Fig. S3b), in our experiments, we saw 2.6-fold increase in ROS in TauT^-/-^ MSCs compared to wild-type controls (Fig. 5g, h).

The elevated ROS levels and defects in oxidative phosphorylation in TauT^-/-^ MSCs were associated with a 2.4-fold increase in cellular senescence in undifferentiated murine TauT^-/-^ MSCs measured by senescence-associated β-galactosidase (SA-β-gal) staining compared to wild-type controls (Fig. 5i, j). Consistent with an induction of senescence in murine MSCs with TauT loss, we also identified a 2 to 4-fold increase in SA-β-gal staining in primary human donor bone marrow derived MSCs expressing shRNAs against *SLC6A6* as compared to a control shRNA (Fig. 5k, l).

Collectively, our data identifies a key role of Wnt/beta-catenin signaling and oxidative phosphorylation downstream of taurine in regulating MSC function (Fig. 5m).

## DISCUSSION

Our work identifies the taurine transporter Slc6a6 as a key regulator of MSC maintenance and function. Our data indicates that the role of taurine in sustaining bone health is possibly downstream of its function in promoting MSC osteogenic fate. Earlier work using osteoblast cell lines has identified Slc6a6 expression in cell culture conditions (30–35). However, our analysis of stromal cells derived from murine bone and bone marrow, as well as osteogenic differentiation of primary bone marrow derived MSCs, indicate that Slc6a6 expression is largely restricted to the MSC and is rapidly downregulated with differentiation. Importantly, we do not detect the expression of other known taurine transporters in our analysis. These data show that TauT expression is restricted to immature MSC population in the murine bone and bone marrow, and that TauT is likely the primary method of taurine uptake by MSCs.

Using a genetic TauT knockout murine model, we show that taurine uptake is essential for sustaining functional MSC populations in vivo. Our data suggests that while taurine uptake is required to promote osteogenic fate of human and murine MSCs, it may be dispensable for adipogenic differentiation. Mechanistically, we find that taurine can impact Wnt signaling and oxidative phosphorylation, both key regulators of MSC differentiation (55). However, we did not observe alterations in adipogenic differentiation with TauT loss. While impaired oxidative phosphorylation can inhibit adipogenesis, increased inflammation and ROS promote it (56, 57), possibly balancing each other and thus leading to no impact on adipogenic differentiation with TauT loss. Consistent with a key role of taurine in sustaining MSC function in the bone marrow, we find that blocking taurine uptake impairs the ability of MSCs to support HSPCs *in vitro*. This is possibly due to the downregulation of growth factors critical for HSPC cell renewal like stem cell factor (SCF) and VEGF-C (58) in the absence of TauT, as seen in our RNA-Seq. Collectively, our data shows that TauT is a key regulator of MSC function and suggests that modulating taurine uptake in these cells may be of therapeutic interest in instances of bone fracture or osteoporosis.

Our data indicates that 16-week-old TauT^-/-^ mice have fewer MSCs and impaired physical and mechanical bone properties *in vivo* as compared to wild-type controls. However, we did not find any changes in muscle morphology or function, indicating that the impact on bone may not be a consequence of declining muscle function at this age. Since MSCs can differentiate into chondrocytes, it may be of interest in future studies to determine the ability of TauT^-/-^ MSCs to differentiate along the chondrocyte lineage. Raman spectroscopy can also be used to test if blocking taurine uptake in MSCs impairs bone mineral composition/structure as the mice age, which may contribute to skeletal abnormalities via disrupted chondrogenesis and osteogenesis with aging.

We find that taurine uptake is critical for maintaining the bone health of young adult mice. The observed osteopenia-like phenotype indicates that it would be of interest to determine the impact of TauT loss on bone fracture repair in these mice. *In vivo*, MSCs play a critical role in the continuous renewal of bone. Notably, MSCs from juvenile mice (under 8 weeks of age) generally exhibit a faster regenerative capacity compared to adult mice (59, 60). It is thus possible that the more pronounced phenotype observed in our work in mice older than 8 weeks results from a decline in MSC regenerative potential, thereby exacerbating the effects of impaired taurine uptake on physical and mechanical bone properties. It is also possible that other signaling molecules may be more crucial during juvenile stages. During early bone development, growth factors like TGF-β and FGF are directly involved in proliferation and differentiation of skeletal system components (61–63). Thus, taurine may not exhibit an effect during this period and could become increasingly important for maintaining bone homeostasis in young adults. The mice used in the scRNA-seq datasets analyzed here (30, 33, 35) were between 6-22 weeks old, thus assessing Slc6a6 expression at earlier developmental stages may be important as its expression may vary during development, and in turn influence bone formation. Future studies would also benefit from the generation of Slc6a6 floxed mice to conclusively establish the temporal requirement of Slc6a6 expression in MSCs on bone development, maintenance, and repair.

The observed differences in bone mineral density (BMD) between males and females are possibly due to distinct hormonal profiles and their effects on the bone remodeling processes. Since Slc6a6 is highly expressed in ovary precursor cells (64), it is possible that Slc6a6 loss in our global knockout mice may be impacting ovarian function. Since ovaries are the primary source of estrogen, and its loss is known to exacerbate bone defects (65), it is possible that differences in estrogen levels may be contributing to the observed variations in BMD between male and female mice. Thus, future studies would benefit from a careful examination of ovarian defects, as well as changes in estrogen levels in TauT^-/-^ mice to determine how estrogen and taurine can work together to impact bone development.

Our observed *in vivo* impact of taurine transporter loss on bone health appears to be less pronounced than the striking effect of TauT loss on MSC differentiation *in vitro*. This could be due to the activation of compensatory mechanisms *in vivo*. Taurine functions as an antioxidant, and impaired taurine transport may promote an inflammatory response. This, in turn, could promote the upregulation of other amino acids that drive anti-inflammatory responses, such as glutamine and arginine, and thus support bone health (66, 67). Upregulation of these amino acids may counteract the detrimental effects of taurine uptake loss on bone health. To conclusively test this, it would be of interest to develop and test methods of metabolic profiling of MSCs directly isolated from the bone and bone marrow.

Importantly, patients with acute myeloid leukemia (AML), or those undergoing chemotherapy for solid tumors often develop osteopenia or osteoporosis and reduced bone density (68–70). We have recently identified a key role of taurine uptake by TauT expressing AML cells in the bone marrow in promoting glycolysis and AML progression (30). It is thus possible that expanding cancer cells deplete available taurine in the bone marrow, leading to reduced taurine availability for MSCs. This may in turn impair bone marrow MSC function and thus contribute to defects in osteogenic differentiation and subsequent osteopenia. Further, we have shown that Slc6a6 expression in MSCs decreases during leukemia progression (30). Thus, Slc6a6 loss may also account for the observed defects in osteogenic differentiation and osteopenia in the context of AML. Given the dual requirement for taurine by leukemia cells as well as MSCs in the bone marrow, it may be of interest to study the dynamics of taurine uptake, and the impact of taurine inhibitors or supplemental taurine on bone health in models of these aggressive leukemias.

## ACKNOWLEDGMENTS

We are grateful to Ember Johnson and Amanda R. Streeter for their technical support. We thank the Cytometry and Genomics Shared Resources supported in part by University of Rochester Wilmot Cancer Institute Support Grant P30CA272302. We are grateful to URMC BMTI and HBMI facilities supported by P30AR069655, and the Mouse Behavior Testing Core. C.M.K. was supported by NIH Training Grant T32AR076950. L.M.C., J.L.L. and R.A.E. are supported in part by NIH grants R01AG079556 awarded to L.M.C. and J.L.L., R01AG076786 to L.M.C. and R.A.E., and R01AG080188 to R.A.E. This work is supported by NIH grants R01DK133131 and R01CA266617 awarded to J.B.

## AUTHOR CONTRIBUTIONS

C.M.K performed majority of the experiments and wrote the paper. S.S. assisted with the biochemical analyses. B.J.R helped with *in vivo* experiments. C.D.B. performed all bioinformatic analyses. P.S. helped with cell culture and histological analysis. T.I. provided TauT knockout mice. K.P.J performed biomechanical testing. C.Y., E.I.F. and E.R.Q. provided experimental assistance. F.C. helped process human donor MSCs provided by J.L.L.. L.M.C, H.A.A, and R.A.E. shared reagents, expertise, and provided intellectual input. J.B. conceived of the project, planned and guided the research and wrote the paper.

## ETHICS DECLARATIONS

### Ethics

All animal experiments were performed according to protocols approved by the University of Rochester’s Committee on Animal Resources. Human samples were obtained from healthy donors after written informed consent in accordance with the Declaration of Helsinki and approval of University of Rochester institutional review board (IRB).

## METHODS

### Analysis of single-cell RNA-sequencing datasets

Data from GEO accessions GSE226644, GSE108892, GSE122467, and GSE128423 was merged and integrated using Seurat v4.1.0 and harmony v0.1.0 within R v4.1.1. Clusters were generated using a resolution of 0.2, marker genes were determined using only positively significantly differentially expressed genes via the FindAllMarkers function, and putative cell types assigned via EnrichR v3.0 against the Azimuth_Cell_Types_2021 database. Clusters expressing CD45, CD71, and Ter119 or annotated as platelets, cycling, or microglia were removed, and this process was repeated.

### Experimental murine model

The *Slc6a6* (TauT) mice were bred as described earlier and are of C57BL/6J background (39). All mice were 4–38 weeks of age and TauT^+/+^ mice were used as controls. C57BL/6J were obtained from Jackson Laboratory. Mice were bred and maintained in the animal care facilities at the University of Rochester. All animal experiments were performed according to protocols approved by the University of Rochester’s Committee on Animal Resources.

### Murine MSC isolation, osteoinduction, and adipoinduction

Primary bone marrow cells were harvested from femurs and tibia bone marrow from TauT^+/+^ and TauT^-/-^ mice as previously described (30, 71). MSCs were plated at a density of 20×10^6^ cells per 10 cm dish coated with 5ug/cm^2^ Type I collagen (Corning). Cells were cultured in MEM α without ascorbic acid (Gibco) supplemented with 15% FBS and 100 IU/mL Penicillin/Streptomycin (Gibco) and incubated at 37°C, 5% CO_2_, and 2% O_2_. Media was exchanged for 3-5 days to remove non-adherent hematopoietic populations. Once cells reached confluency (7-10 days), MSCs were enriched using magnetic depletion (CD45^-^Ter119^-^CD31^-^) or by flow sorting. Purified MSCs were plated at a density of 20×10^3^ cells/cm^2^ in a 10 cm plate coated with 5ug/cm^2^ Type I collagen (Corning). Cells were osteoinduced at confluency in their media consisting of MEM α (Gibco) supplemented with 10% FBS, 100 IU/mL Penicillin/Streptomycin (Gibco), 50 mg/ml ascorbic acid (Sigma-Aldrich), and 2.5 mM β-glycerolphosphate (Sigma-Aldrich) for 14 days. Cells were adipoinduced at confluency in their media consisting of MEM α without ascorbic acid (Gibco) supplemented 10% FBS, 100 IU/mL Penicillin/Streptomycin (Gibco), 1 µM Dexamethasone (Cayman Chemical), 0.5 mM IBMX (Sigma), 10 µg/ml Insulin (Humulin) and 1 µM Rosiglidazone (Cayman Chemical). Cells were incubated in induction media for 2 days, then changed to maintenance media consisting of MEM α without ascorbic acid (Gibco) supplemented 10% FBS, 100 IU/mL Penicillin/Streptomycin (Gibco), 10 µg/ml Insulin (Humulin) and 1 µM Rosiglidazone (Cayman Chemical) for 7 days. Cells were stained for mineralization using Alizarin Red S or alkaline phosphatase and Von Kossa and quantified as previously described (72–74). Cells were stained for lipid formation using Oil Red O and quantified as previously described (41, 74, 75).

### RNA extraction and qRT-PCR

RNA was extracted using the RNeasy Micro kit (Qiagen) as per the manufacturer’s protocols. RNA concentrations were determined using NanoDrop 1000 Spectrophotometer (Thermofisher Scientific). RNA quality was assessed with the Agilent Bioanalyser 2100 (Agilent Technologies). qRT-PCR was carried out on BioRad CFX96 C100 Thermocycler using BioRad CFX Manager 1.1 v4.1 (BioRad) or Thermofisher Scientific Quant Studio 12K Flex Real Time PCR using Quant Studio v1.2 (Thermofisher Scientific). qRT-PCR data was analysed using BioRad CFX Manager 1.1 v4.1 or Quant Studio v1.2.

### Stromal cell isolation and FACS analysis

Microenvironmental populations were isolated as previously described (30, 76). Briefly, bone and bone marrow (BM) cells were isolated from long bones in 1x HBSS (Gibco) with 5% FBS (GeminiBio) and 0.5 M EDTA (Gibco). BM cells were digested for 30 minutes in 1x HBSS containing 2 mg/mL Dispase II (Gibco), 1 mg/mL Collagenase Type IV (Sigma-Aldrich), and 20 ng/mL DNase Type II (Sigma-Aldrich). Bone spicules were digested for 45 minutes in 1x PBS supplemented with 2.5 mg/mL Collagenase Type I (Stem Cell Technologies) and 20% FBS. Digested BM was RBC lysed using RBC Lysis Buffer (eBioscience). Bone and BM cells were pooled and CD45^+^ Ter119^+^ hematopoietic cells were magnetically depleted on an autoMACS cell separator (Miltenyi Biotec). The CD45^-^Ter119^-^ stromal cells were stained and analysed for candidate populations by flow cytometry on LSRFortessa (BD Biosciences). Antibodies used for defining stromal cell populations were as follows: CD45^-^Ter119^-^CD31^+^Sca1^+^ (arteriolar endothelial cells), CD45^-^Ter119^-^ CD31^+^Sca1^-^ (sinusoidal endothelial cells), CD45^-^Ter119^-^CD31^-^Sca1^+^CD51^+^ (MSCs). Data was analyzed using FlowJo software.

### Fibroblast (CFU-F) and osteoblast (CFU-OB) colony formation assays

Bone marrow cells were isolated from mice as described above. Bone marrow cells were seeded at a density of 25,000 cells/cm^2^ in 6-well plates. Media was exchanged for 3 days to remove non-adherent hematopoietic populations. Osteoinduction was performed as described above. For assessment of CFU-F, cells were cultured in MEM α without ascorbic acid (Gibco) supplemented with 15% FBS (GeminiBio) and 100 IU/mL Penicillin/Streptomycin (Gibco). Both CFU-OB and CFU-F were incubated at 37°C, 5% CO_2_, and 2% O_2_ for 14 days. On day 14, cells were either stained with Crystal Violet (CFU-F) or ALP and Von Kossa (CFU-OB) to assess total colonies. Clusters of greater than 50 cells were considered a CFU. Plates were imaged on a LI-COR Odyssey M. Cells were quantified using ImageJ.

### Primary human donor bone marrow derived MSC osteogenic differentiation

Primary human donor bone marrow MSCs were cultured from marrow aspirates from healthy donors after written informed consent in accordance with the Declaration of Helsinki and approval of University of Rochester institutional review board (IRB). Osteogenic differentiation was carried out as described above. All shRNA sequences are as described earlier (30).

### Coculture of KLS cells and MSCs

MSCs were isolated from TauT^+/+^ and TauT^-/-^ mice as stated above. 100,000 MSCs per well were plated in a 24-well plate. After 24 hours, ∼20,000 C57BL/6J KLS cells (Lin-cKit+Sca1+) were FACS sorted and cocultured with MSCs in RPMI (Gibco) media supplemented with 10% Hi-FBS (GeminiBio), 50 mM 2-mercatpoethanol, and Penicillin-Streptomycin (Gibco), media adapted from previous publication (77). Post 72 hours, the hematopoietic suspended cells were removed for viability analysis, FACS, and CFU assays. Viability was assessed using 0.4% Trypan blue (Thermofisher) on an Olympus CK2 microscope. Cell lineage content was assessed via FACS on LSRFortessa (BD Biosciences). Antibodies used for defining lineage^+^ are as followed: CD3ε, CD4, CD8, Gr1, CD11b/Mac-1, Ter119, CD45R/B220 and CD19. Data was analyzed using FlowJo software. Colony forming ability was assessed by plating 400 cells in methylcellulose medium (M3434, StemCell Technologies). Colonies were scored on day 7 using an Olympus CK2 microscope.

### Dual-energy X-ray absorptiometry (DEXA)

DEXA scans were carried out using equipment at the University of Rochester Biomechanics and Multimodal Tissue Imaging (BMTI) Core. Male and female TauT^+/+^ and TauT^-/-^ mice were measured between 4 and 38 weeks of age. Mice were weighed, then anesthetized using either 100 mg/kg Ketamine and 10 mg/kg Xylazine IP at 0.1 mL per 10 grams of body weight or 2% isoflurane vapor. Under anesthesia, the mouse’s whole body was scanned, and an area of interest was selected using Lunar PIXImus2 system. The whole body and femur were analyzed in terms of bone mineral density (BMD), bone mineral content (BMC) and fat percentage.

### Bone micro-computed tomography (micro-CT)

Micro-CT was completed at the University of Rochester BMTI Core. TauT^+/+^ and TauT^-/-^ femurs were isolated, cleaned of excess soft tissue, and stored in 1x PBS at-80°C prior to micro-CT. Specimens were imaged using high-resolution acquisition (10.5 µm voxel size) with the VivaCT 40 tomograph (Scanco Medical). Scanco analysis software was utilized for volume quantification.

### Biomechanical torsional testing

Biomechanical torsional testing was completed in collaboration with the University of Rochester BMTI Core. TauT^+/+^ and TauT^-/-^ femurs were isolated, cleaned of excess soft tissue, and stored in 1x PBS at-80°C prior to biomechanical testing. Tibia samples were held in bone cement and tested using EnduraTec TestBench system (Bose). The tibiae were tested in torsion until failure at a rate of 1°/s. The torque data were plotted against rotational deformation to determine maximum torque and torsional rigidity.

### Grip strength test

Grip strength testing was completed at the University of Rochester Mouse Behavior Core according to previously published protocols (78). Grip strength was analyzed for limbs using a digital force meter (Columbus Instruments Grip Strength Meter) equipped with precision force gauges to retain the peak force applied. Each mouse was tested 5 times on the limbs with a minimum 60 second inter-trial interval.

### Histological staining

Histology staining was completed at the University of Rochester Histology, Biochemistry, and Molecular Imaging (HBMI) Core. TauT^+/+^ and TauT^-/-^ femurs were isolated, cleaned of excess soft tissue, and were fixed for three days in 10% neutral buffered formalin (NBF) solution and processed for histology via decalcification in Webb-Jee 14% EDTA solution for two weeks followed by paraffin embedding. Samples were sectioned at 5 µm in three levels of each sample and then stained with tartrate-resistant acid phosphatase (TRAP). TauT^+/+^ and TauT^-/-^ gastrocnemius muscles were isolated, cleaned of excess soft tissue, and were fixed for two days in 10% NBF solution and processed for histology. Samples were sectioned at 5 µm in three levels of each sample and then stained with hematoxylin and eosin (H&E). Microscopy images were obtained on Olympus CKX41 using CellSens Entry v2.3 (Olympus). The number of TRAP positive cells and myofiber cross-sectional area was measured in ImageJ.

### Flow cytometry-based analysis of CellROX

MSCs were isolated from TauT^+/+^ and TauT^-/-^ mice as stated above. CellROX (ThermoFisher) was added to 50,000 MSCs at a final concentration of 20 µM in MEM α without ascorbic acid (Gibco) supplemented with 15% FBS (GeminiBio) and 100 IU/mL Penicillin/Streptomycin (Gibco). Cells were incubated with CellROX for 30 minutes at 37°C. Cells were washed three times with 1x PBS, and cells were analyzed via FACS on LSRFortessa (BD Biosciences). Mean fluorescence intensity of CellROX was quantified using FlowJo.

### *Bulk RNA-sequencing of* TauT^+/+^ and TauT^-/-^ MSCs

1ng of total RNA was used as input in the SMART-Seq mRNA LP (Takara Bio USA, San Jose, CA) library prep workflow per manufacturer’s recommendations. Briefly, full-length cDNA was synthesized via oligo-dT priming and template-switching at the 5’ end. cDNA libraries were then amplified by long-distance PCR. The quantity and quality of the amplified cDNA was determined using the Qubit Fluorometer (Life Technologies, Carlsbad, CA) and the Agilent Bioanalyzer 2100 (Santa Clara, CA). Subsequently, 1ng of amplified cDNA was used for library construction through enzymatic fragmentation and ligation of stem-loop adapters, which provide binding sites for final library amplification. Lastly, Illumina-compatible libraries were amplified using unique dual-indexed primers. Amplified libraries were assessed for quantity and quality using the Qubit Fluorometer and the Agilent Fragment Analyzer and prepared to sequence on the Illumina NovaSeq X Plus (Illumina, San Diego, CA) 10B flowcell with paired-end reads of 150nt.

Raw reads generated from the Illumina basecalls were demultiplexed using bcl-convert v4.1.7. Quality filtering and adapter removal was performed using FastP v.0.23.1. Processed reads were then mapped to the human reference genome (GRCm39 + gencode M31) (https://www.gencodegenes.org/mouse/release_M31.html) using STAR_2.7.9a. Reads mapping to genes were counted using subread featurecounts v2.0.1 with “-s 2”. Differential expression analysis was performed using DESeq2-1.34.0 with an adjusted P-value threshold of 0.05 within R v4.1.1 A design formula incorporating both sex and condition was used to regress out the impacts of sex in differential expression. Gene ontology analyses were performed using the EnrichR-3.0 package.

### Western blot analysis

MSCs were isolated from TauT^+/+^ and TauT^-/-^ mice as described above. Cell lysates were prepared in 1x RIPA (Thermo Scientific) supplemented with 1x protease/phosphatase inhibitors (Cell Signaling Technology, CST) and 250IU Benzonase Nuclease (Millipore Sigma). Samples were separated on gradient polyacrylamide gels and transferred to nitrocellulose blotting membrane (0.45µM; GE healthcare). Primary antibodies against non-phosphorylated active beta-catenin (β-catenin) and β-actin (CST) were used. Horse-radish peroxidase (HRP)-conjugated anti-rabbit antibody (CST) was used to detect primary antibodies. Immunoblots were developed using SuperSignal West Femto Maximum Sensitivity Substrate (Thermo Scientific). Immunoblots were imaged using LI-COR Odyssey M using Empiria Studio v2.3 (LI-COR). Images were analyzed using Empiria Studio v3.2.0.186 (LI-COR).

### Seahorse assays

Seahorse XF cell mito stress test kit (Agilent Technologies, 103015-100) was used to measure glycolytic flux (ECAR) and oxygen consumption (OCR) respectively. 20,000 MSCs were seeded in 96-well XF96 well plates in MEM α without ascorbic acid (Gibco) supplemented with 15% FBS (GeminiBio) and 100 IU/mL Penicillin/Streptomycin (Gibco). Cells were incubated for 24 hours at 37°C, 5% CO_2_, and 2% O_2_. Following 24 hours, OCR and ECAR data was measured following sequential addition of oligomycin (1.5µM), carbonyl cyanide-4 (trifluoromethoxy) phenylhydrazone (FCCP) (0.5µM), rotenone and antimycin (0.5µM) using XF96 analyzer. Data was analyzed using Wave v2.6.3 (Agilent Technologies).

### Senescence associated beta-galactosidase staining of murine MSC

Human and murine MSCs were cultured as described above. Senescence associated beta-galactosidase staining was completed according to manufacturer’s protocol (Cell Signaling Technology). Cells were analyzed using ImageJ.

### Statistical analysis

Statistical analyses were carried out using Graphpad Prism software v6.0 (GraphPad software Inc.). Data are mean ± SEM, mean ± SD. One-way ANOVA, unpaired two-sided Student’s t-tests, and multiple unpaired t-tests corrected with Benjamin & Hochberg method, were used to determine statistical significance.

## SUPPULMENTARY FIGURE LEGENDS

**Figure S1:**
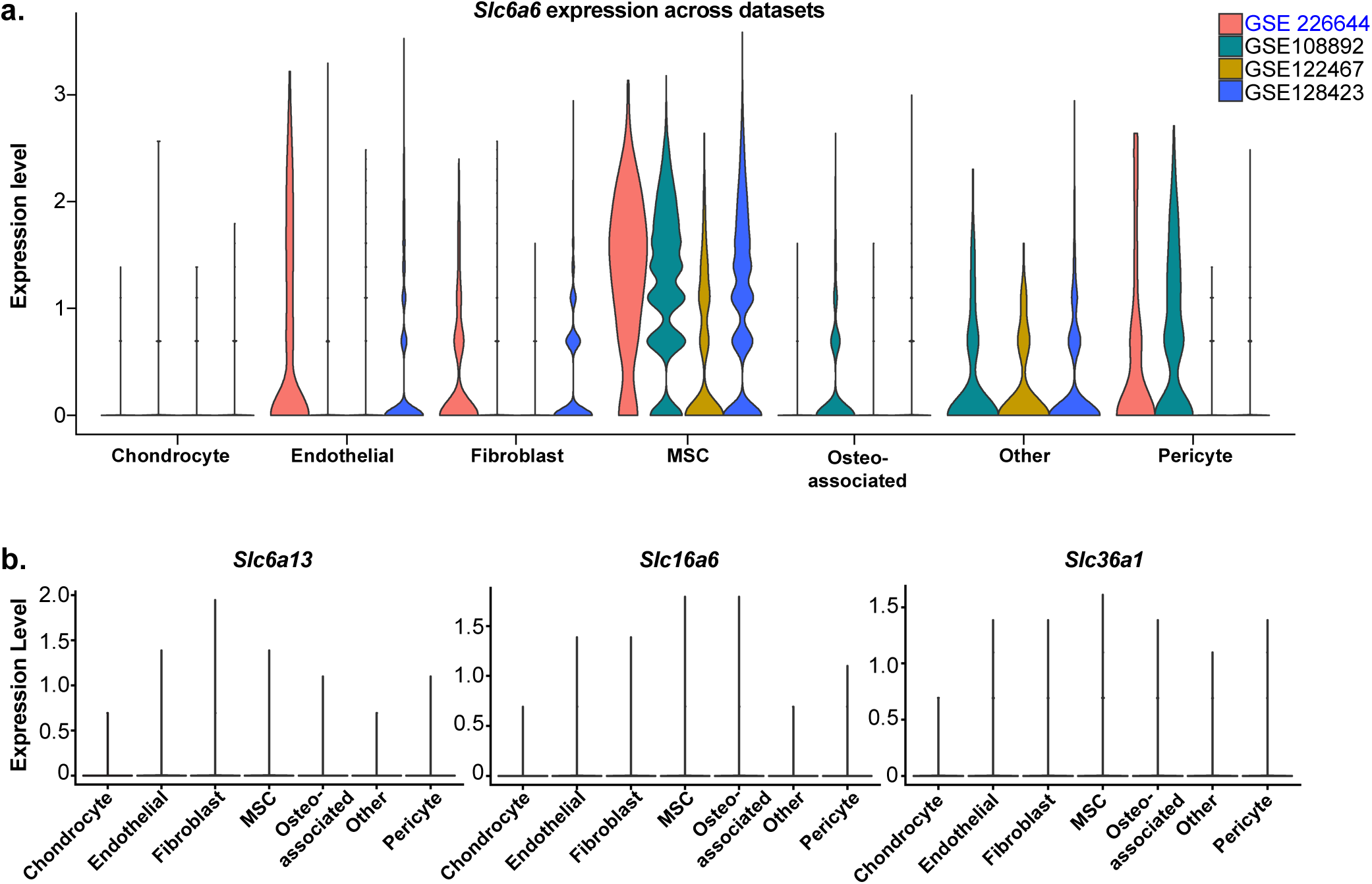
Mesenchymal stromal cells have the highest expression of Slc6a6. **a, b,** Violin plots of Slc6a6a expression (**a**) and Slc6a13, Slc16a6, and Slc36a1 expression (**b**) in non-hematopoietic cells in publicly available scRNA-sequencing data sets (blue text indicates our sc-RNA-sequencing data set).

**Figure S2:**
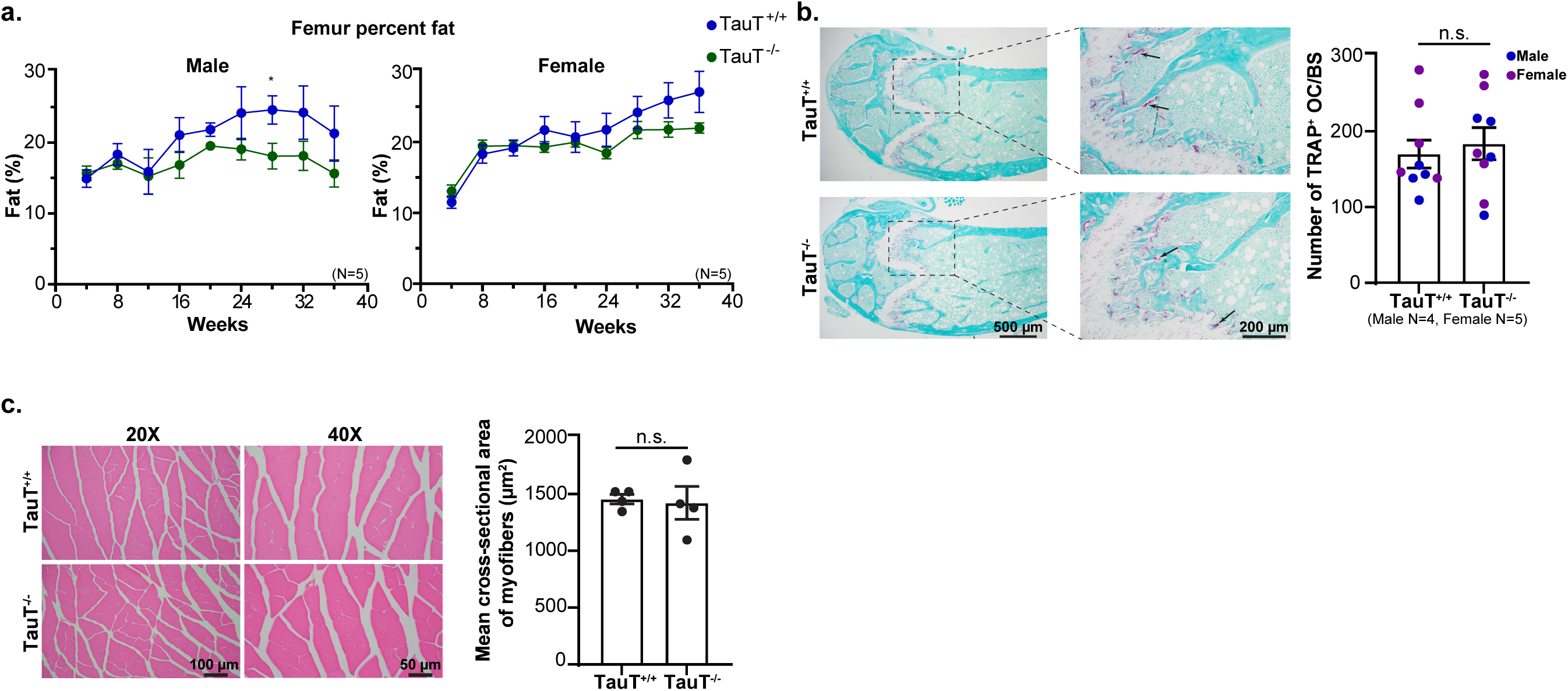
MSCs are responsible for observed physical bone defects in the absence of TauT *in vivo.* **a**, DEXA scans of femur percent fat of TauT^+/+^ and TauT^-/-^ mice over a 40-week period (mean ±s.e.m.; data are combined from four independent experiments). **b**, Representative tartrate resistant acid phosphatase (TRAP) staining of murine femurs and quantification (mean ±s.e.m.; data are combined from four independent experiments). **c**, Representative Hematoxylin and Eosin (H&E) staining of gastrocnemius muscle and quantification of myofiber cross-sectional area (mean ±s.e.m.). (*p<0.05, **p<0.01. ***p<0.001). All analyses are from unpaired two-tailed Student’s t-test.

**Figure S3:**
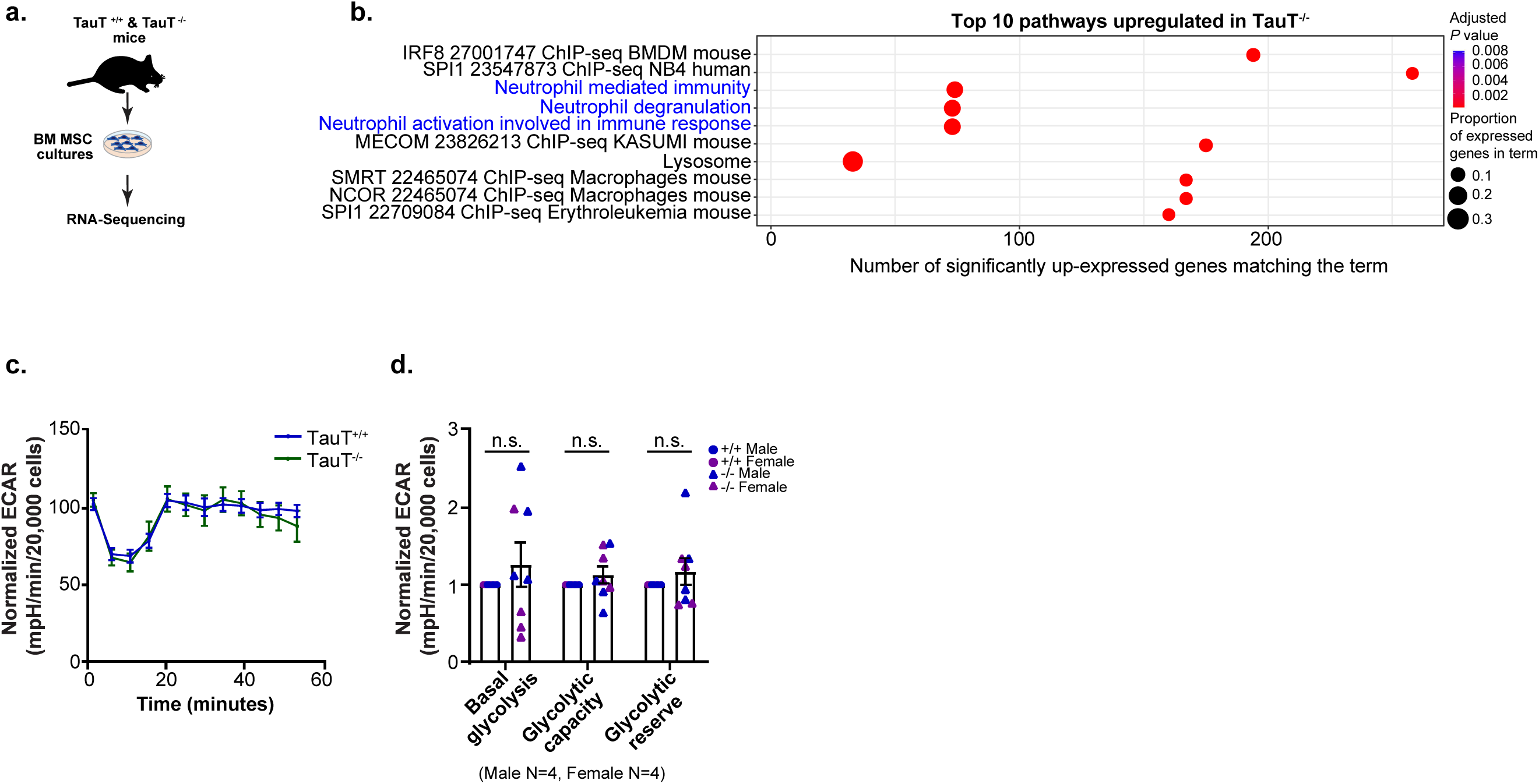
Increased inflammation may further contribute to MSC osteogenic defects in the absence of TauT. **a**, Experimental strategy used for bulk-RNA sequencing of male and female TauT^+/+^ and TauT^-/-^ MSCs (n=8 male and n=7 female MSCs). **b**, Gene sets significantly upregulated in TauT^-/-^ MSCs (blue text indicates pathways of interest). **c**, **d**, Extracellular acidification (ECAR) curve (**c**) and normalized quantification (**d**) (mean±s.e.m.; n=4-5 independent culture wells per cohort; data combined from four independent experiments; unpaired two-tailed Student’s t-test).

